# Can root systems redistribute soil water to mitigate the effects of drought?

**DOI:** 10.1101/2022.09.15.508112

**Authors:** Andrew Mair, Lionel X Dupuy, Mariya Ptashnyk

## Abstract

Plants adapt both morphologically and physiologically in response to drought. This work considers another potential mechanism of plant drought resistance. Namely, the capacity of root systems to modify the hydraulic properties of surrounding soil, and extend the post-precipitation lifetime of water in the rooted zone.

In studies of plant drought resistance, the effect of root-oriented preferential soil water transport is seldom considered, despite experimental evidence that it occurs. We developed a model for water transport through soil that incorporates preferential flow induced by a root system. Bayesian optimisation was employed to calibrate the model against experimental data, and the finite element method was used to obtain model simulations for the influence of different root architectures on post-precipitation water lifetime.

When increasing preferential flow strength, simulations indicated a trade-off between evaporation and deep percolation losses. It was also observed that the soil surrounding root systems with a reduced gravitropic response, retained the most water following precipitation.

This work provides new insight into the role of root system traits in plant drought resistance, and identifies potential crop phenotypes for improved water use efficiency.

## Introduction

Drought resistance in crops is due to the ability of plants to avoid dehydration through morphological and physiological mechanisms (Fang and Xiong, 2015). Primary responses usually involve adapting the processes that govern transpiration. For example, plant species regulate the opening and closure of their stomata in order to adjust levels of water loss via transpiration (Luo, 2010). Leaves can also produce waxes (Goodwin and Jenks, 2005), and roll (Cal et al.,2019), to reduce permeability and transpiration area respectively.

A number of physiological root characteristics have been linked to improved drought resistance. The development of the root exodermis, by the laying down of Casparian bands and suberin lamellae, improves water retention by altering the radial hydraulic conductivity of root tissue (Frensch and Steudle,1989; Taleisnik et al., 1999; Hose et al., 2001). Plants can also increase solute concentration in their root cells, thus enabling water extraction at lower soil water potentials (Li et al., 2008). The impact of root system architecture on drought resistance has also received much attention. Primarily with the aim of identifying the optimal spatial distribution of roots, so that water in deep soil can be accessed (Uga et al., 2013).

Plant root activity has a profound impact on the hydraulics of vegetated soil. The exudation of mucilage by roots has been shown to modify the water repellency of rhizosphere soil (Ahmed et al., 2016; Naveed et al., 2018). Furthermore, root microbiome activity and the physical forces of growing roots, influence the pore structure of surrounding soil. Several studies, on different soil types, provide evidence of altered porosity near root surfaces (Dexter, 1987;Bruand et al., 1996; Feeney et al., 2006; Koebernick et al., 2019; Anselmucci et al., 2021). Overall, these modifications to soil properties have been found to manifest as an increase in the transport of soil water into directions that follow the orientation of roots (Noguchi et al., 1997; Michot et al., 2003; Lange et al.,2009; Beff et al., 2013). This is described as preferential flow (Ghestem et al.,2011), and is likely to influence the fate of all water that enters vegetated soil.

Previous research on drought resistance has not greatly considered the ability of plants to reduce soil water loss through the mechanism of preferential flow. Prior to canopy closure, evaporation from the soil surface is a major source of water loss from vegetated soil (Schwinning and Sala, 2004). However, the leaching of water into soil layers below the root profile, referred to as deep percolation, also contributes to total water loss (Bethune et al., 2008). As does surface run off, where precipitation fails to infiltrate into bulk soil due to the surface already being fully saturated. Water is also removed from the soil via root uptake and either used for cell expansion and growth, or released back to the atmosphere via transpiration. Generally, most water that enters a soil is lost to the environment instead of being used by plants. This is illustrated by the fact that plants growing in hydroponic systems have been found to require 80 to 90 percent less irrigation than when grown in soil (AlShrouf et al., 2017).

This work aims to investigate how the architecture of a root system influences the way that soil water is redistributed following precipitation. The directions of water transport in the soil determine not only its availability for uptake by roots, but also the rate at which it is lost from the rooted soil layer. We therefore hypothesise that root system architecture can affect the post-precipitation water lifetime, which is the length of time that water remains available for plant uptake (Figure 1). A model is proposed for water transport through vegetated soil, which combines root system architecture, root water uptake (RWU) and root-oriented preferential flow (Mair et al., 2022). Our model was calibrated for Maize plants with respect to experimental data, on the hydraulic conductivity of soil vegetated by Maize (Feki et al., 2018). Simulations were performed to investigate the effect of root-oriented preferential flow on evaporation, deep percolation, and root water uptake. We examined how our results changed depending on the architecture of a root system, and hence proposed ideotypes for improved drought resistance.

**Fig. 1:**
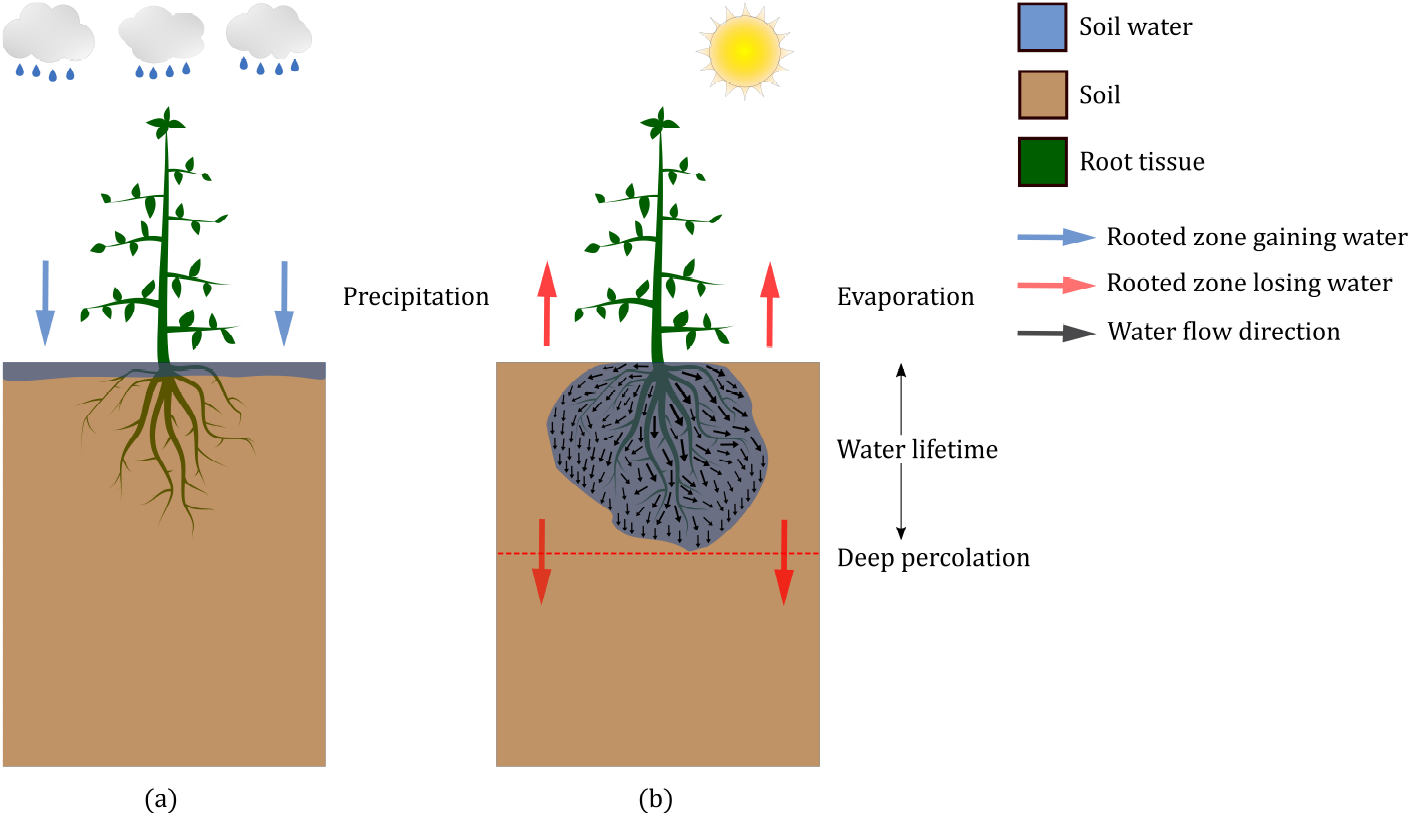
Root system effects on soil hydraulics and drought resistance. Drought occurs when a water deficit is maintained for sufficiently long enough for plant dehydration to occur. Following precipitation (a), root systems redistribute soil water. The spatial distribution of water then affects evaporation and deep percolation rates (b), and hence the duration of any water deficit imposed on the plant.

## Materials and Methods

### A model for soil water transport incorporating root uptake and preferential flow

This work considers a root-soil system of a 3D domain of unsaturated soil Ω occupied by a root system 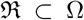. Our model for water transport in the vegetated soil incorporates the influence of root system architecture by using density functions, which give continuous approximations of root abundance and orientation within the soil. The root system is defined as a finite union of 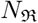 disjoint segments 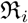, i.e. 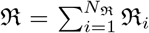. Each root segment is defined by values for the positions and radii of its base and tip. Such data is obtained for real root systems through excavation and digitisation (Danjon et al., 1999), or the use of x-ray computed tomography Zhao et al. (2020). It is also the typical output of root system architecture simulation platforms such as CRoot-Box (Schnepf et al., 2018) and OpenSimRoot (Postma et al., 2017). In this work the software CRootBox is used, which employs an L-system model (Leitner et al., 2010) to create virtual root system architectures, for a number of species, from parameters such as root length, inter-lateral distance and gravitropism.

Using the methods in Mair et al. (2022), we derive, from architecture data of a root system, the volumetric root density *ψ* and the root length density *RLD*. In short, for a spatial co-ordinate *x* in the soil Ω, the value of *ψ*(*x*) ∈ (0,1) increases with root volume, and the value of *RLD* (*x*) increases with root length. Furthermore, *ψ* and *RLD* integrate over the vegetated domain to give the total volume and root length of the root system respectively. Root-oriented preferential flow is incorporated into the model for water transport by *ψ*, and *RLD* is used to model root water uptake.

To model the influence of root architecture and abundance on the transport of water through soil, we use the recently developed model in (Mair et al.,2022),

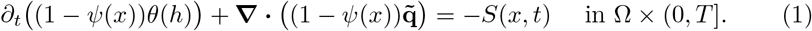

This is a modification of Richards equation, (Richards, 1931), the classic model for the evolution of water content *θ* (L^3^L^-3^) within unsaturated soil, and is defined over a domain 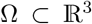, with final time *T* > 0. Equation (1) is solved for pressure head *h* (L). Root water uptake is accounted for by the sink term *S* (T^-1^), and, to incorporate the phenomena of root-oriented preferential flow, the water flux used in (1) has the following form:

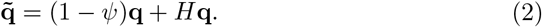

Here **q**(LT^-1^) is the isotropic flux for water flow through fallow unsaturated soil. Root-oriented preferential flow is incorporated by the flow anisotropy matrix *H*, which is parametrised by the facilitation constant *c_a_* > 1 to control the strength of preferential flow induced. The entries of H vary in value over the soil domain according to the location and orientation of roots within the soil. Furthermore, the magnitude of *H* is proportional to *ψ*, meaning that in soil regions with low root abundance the first term in 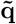 will dominate, and the flux will closely resemble **q**(LT^-1^). However, in regions of soil close to roots the second term in 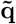 dominates, and the flow of soil water will be facilitated in directions parallel to root axes.

The isotropic flux of water through fallow unsaturated soil is modelled using the Darcy-Buckingham law (Darcy, 1856; Buckingham, 1907):

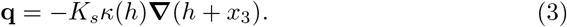

Here, the soil’s saturated hydraulic conductivity is given by the constant *K_s_*(LT^-1^), and the relation between hydraulic conductivity and pressure head *h* is described by the function *κ*, defined using the classic model of Van Genuchten (1980):

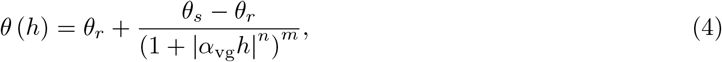

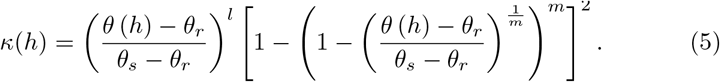

Parameter values in (4) and (5) depend upon soil type. The residual and saturated water contents are given by *θ_r_* and *θ_s_* (L^3^L^-3^) respectively, and α_vg_(L^-1^), *n* (–) and *m* =1 – 1/*n* are shape parameters. The parameter *l* denotes the tortuosity.

Root water uptake is incorporated into the model (1) through the sink term *S*, which comes from existing models (Simunek and Hopmans, 2009;Cai et al., 2018) that link soil water pressure head, normalised root length density *NRLD* (L^-3^), and potential plant transpiration 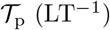:

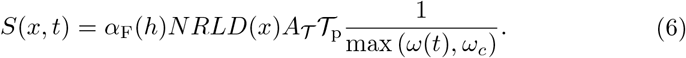

Here 0 ≤ *α*_F_ ≤ 1 is a dimensionless water stress response function that accounts for the impact of reduced water availability on root water uptake (Feddes, 1982). A dependence of root water uptake on the distribution of roots within the soil is incorporated into *S* through the normalised root length density function:

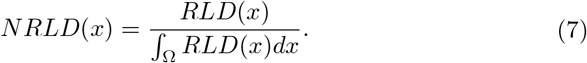

The soil surface area associated with transpiration is denoted as 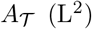, and the dimensionless function *ω*(*t*) is a measure of the global water stress experienced by the plant at a given time (Cai et al., 2018). The value of the critical water stress index *ω*_c_ ∈ (0,1) reflects the plant’s capacity to compensate for low water availability in certain regions of vegetated soil, by increasing uptake in wetter regions.

In this work the soil domain Ω is assumed to be cuboidal. The upper boundary, at the soil-atmosphere interface *x*_3_ = 0, is denoted by Γ_1_, and is where water is lost via evaporation. Furthermore, the seeding location of the root system 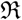, is taken to be (0, 0, 0) ∈ Γ_1_. The lower boundary and system rooting depth, are denoted by Γ_3_, and 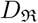 respectively. The lateral boundaries of the domain are denoted by Γ_2_, and water loss through these surfaces is assumed zero. The initial pressure head profile and boundary fluxes are then defined mathematically as

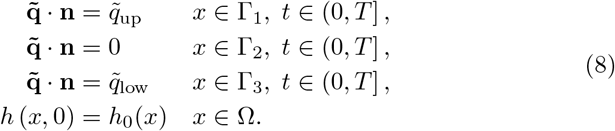

The water flux at the upper boundary is 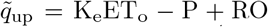, with the precipitation and runoff of water denoted by P and RO (LT^-1^) respectively. Evaporation is K_e_ET_o_ where the function K_e_ determines the proportion of total potential evapotranspiration that comes from evaporation (Allen et al.,1998). We formulate the runoff function as

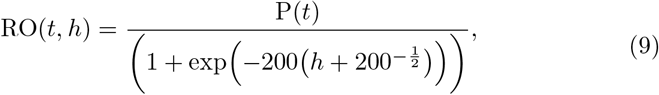

so that if a section of soil on the upper surface is fully saturated, then all precipitation falling onto this section is lost as runoff instead of infiltrating into the soil. At the lower boundary Γ_3_ a free drainage condition is imposed (Rassam et al., 2003):

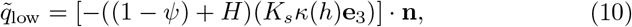

where **e**_3_ is the unit vector in the upward *x_3_* direction. If not provided here, then explicit expressions for the functions discussed in this section are given in the supplementary material.

### Model parametrisation and calibration

The majority of parameters in model (1), (8), were taken from existing literature. Values for *K_s_*, along with the parameters in the hydraulic conductivity (5), and water retention functions (4), came from (Carsel and Parrish, 1988). The tortuosity in (5) was set to *l* = 0.5 (Van Genuchten and Pachepsky, 2011), and parameter values in the functions for uptake and evaporation were taken from (Allen et al., 1998; Cai et al., 2018; Wesseling, 1991). The *Zea mays 1* dataset of CRootBox was used to simulate 5 Maize root systems, each 90 days old, for which density functions and flow-anisotropy matrices were constructed in order to obtain corresponding parametrisations of model (1), (8). These root systems were assumed to be static during the time span of each simulated scenario. The first root system 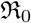 was generated using default growth parameter values for Maize, and used to calibrate model (1), (8) against experimental data. Here the soil domain was assumed to have lateral surfaces at the extremal points of 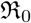, and to reach twice as deep as rooting depth (Figure 2 (a)). The second simulated root system 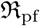, also obtained using default growth parameter values, was used to investigate the influence of preferential flow strength on uptake and water lifetime in the rooted zone. The final 3 simulated root systems were used to investigate the impact of root architectural traits on uptake and water lifetime in the rooted zone. These were generated so that the total volume occupied by each was within a 2% tolerance of the others. One was a control root system 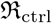, generated using default growth parameter values (Figure 4 (a)). Another, 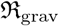, was generated with gravitropism parameters reduced, so that its roots grew further in lateral directions before growing downward (Figure 4 (b))), and a final system 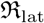 was generated with growth parameter values that gave it similar characteristics to 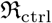, but with longer, and fewer, lateral roots, and slightly shorter primary roots (Figure 4 (c)). Identical soil domains were used for each of the root systems 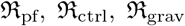, and 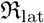, with the lower boundary defined by the depth reached by the deepest root from all four systems. For simulations that considered a silt loam soil, an initial pressure head condition of *h*_0_ = −5m was prescribed, giving an initial water content *θ*(*h*_0_) = 0.214m^3^m^-3^. When considering other soil types, different values were set for *h*_0_ so that *θ*(*h*_0_) = 0.214m^3^m^-3^ was maintained. In the scenario used for calibration, a final time of *T* = 2 days was used and the precipitation condition P_0_ prescribed a single rainfall event on each day. In the simulation for the impact of preferential flow strength on water lifetime we set *T* = 4 days, and in simulations for the influence of root architectural traits we set *T* = 7 days. However, in both cases we used a precipitation condition P that prescribed rainfall only on day 1. Expressions for P_0_ and P are detailed in the supplementary material. Tables 1, 2, and 3 in the supplementary material provide specific sources for all parameters in model (1), (8), along with the values assigned.

**Fig. 2:**
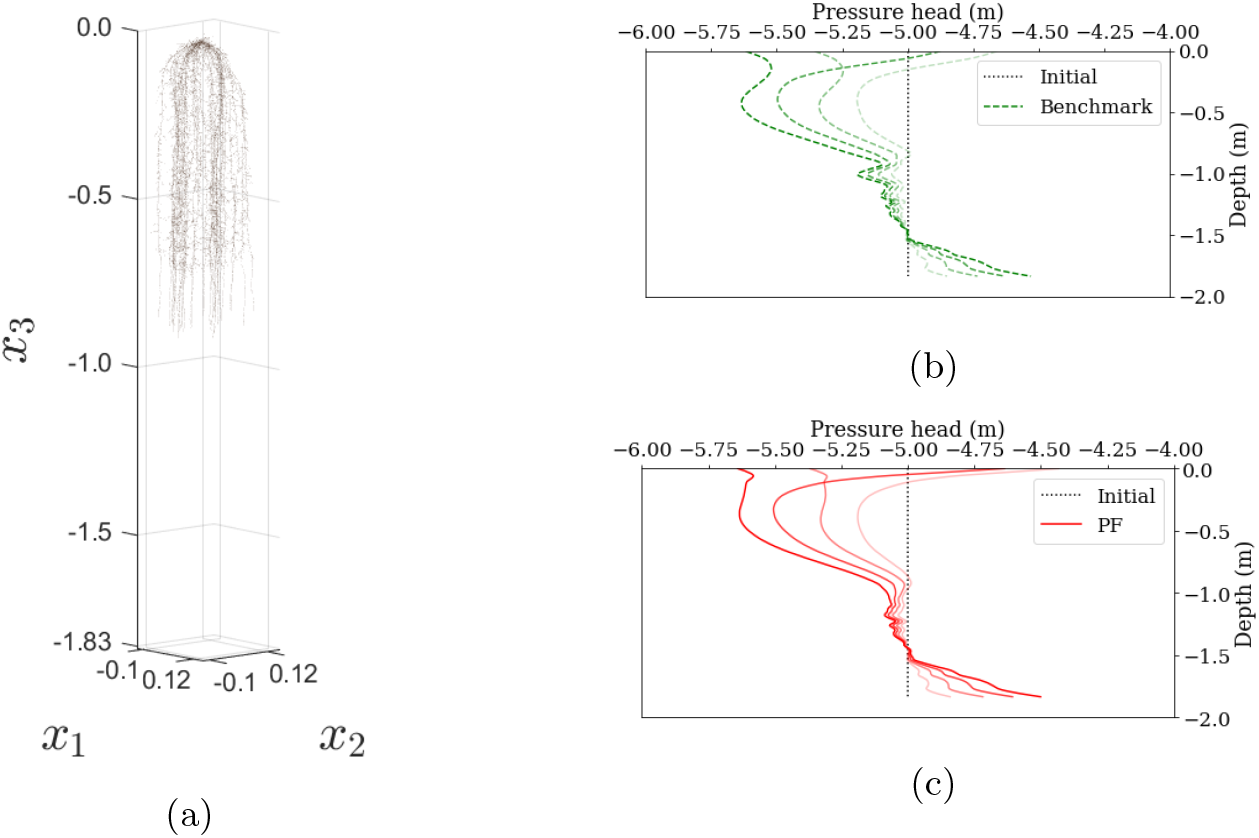
The capacity of model (1), (8) to replicate pressure head profiles from the benchmark model. (a) Simulated Maize root system for which (i) benchmark pressure head profiles were obtained and (ii) the model (1), (8) was parametrised. (b) The benchmark pressure head profiles generated for soil vegetated by the Maize root system shown in (a). (c) The pressure head profiles from model (1), (8), labelled “PF”, when calibrated with the facilitation constant 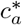 for the Maize root system in (a). In (b) and (c) lines of full opacity show pressure head profiles at the final time (*T* = 2 days), and fainter lines show profiles at earlier times.

**Table 1:** Impact of root system architecture on water lifetime quantities. Results were computed using simulations of water transport from model (1), (8) when parametrised for root systems with different architectural characteristics (Figure 4).

| Root system | Cumulative water lifetime quantities ($\text{m}^3$) | | | |
| --- | --- | --- | --- | --- |
|  | Evaporation | Deep percolation | Total water loss | Water uptake |
| Control $\mathfrak{R}_{\text{ctrl}}$ | $4.43 \times 10^{-3}$ | $9.04 \times 10^{-4}$ | $5.33 \times 10^{-3}$ | $3.26 \times 10^{-2}$ |
| Reduced gravitropism $\mathfrak{R}_{\text{grav}}$ | $4.30 \times 10^{-3}$ | $8.41 \times 10^{-4}$ | $5.15 \times 10^{-3}$ | $4.22 \times 10^{-2}$ |
| Increased lateral length $\mathfrak{R}_{\text{lat}}$ | $4.44 \times 10^{-3}$ | $8.46 \times 10^{-4}$ | $5.29 \times 10^{-3}$ | $3.41 \times 10^{-2}$ |

The only parameter value left to determine in model (1), (8) was the axial facilitation constant *c_a_*. This was achieved through calibration against experimental measurements of Feki et al. (2018), which identify a value 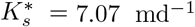, for the saturated hydraulic conductivity of soil, when vegetated by a 90 day old Maize root system. Firstly, Richards equation was parametrised with a depth dependent saturated hydraulic conductivity that took the value 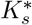 in the vegetated section of soil, and the value for fallow soil in the section below. By adding a sink term for the water uptake of root system 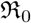, and equipping the equation with boundary and initial conditions equivalent to those in (8), a Benchmark model was formed. Numerical pressure head profiles from this model provided a reference, for the effect of Maize root systems on soil hydraulic properties, against which the value of *c_a_* in model (1) (8) could be calibrated. A cost function *u* was then formulated whose minimizer 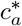 would estimate the facilitation constant value with which to parametrise model (1) (8), so that numerical pressure head profiles accurately matched the reference profiles provided by the benchmark model. Full details on the formulation of *u* and the benchmark model, are given in the supplementary material. A Bayesian optimisation scheme was employed to find the minimiser 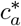 of the cost function. This method was chosen because it uses a probabilistic framework, which requires relatively few cost function evaluations to efficiently explore the parameter space, and does not rely on access to derivatives (Brochu et al., 2010). These are desirable traits because *u* is essentially non-differentiable, and each evaluation at a given *c_a_* value is time-intensive.

### Simulated scenarios

To investigate the influence of preferential flow strength on root water uptake, and water lifetime in the rooted zone, the model (1), (8) was parametrised for the root system 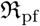, and simulated for 13 different values of facilitation constant *c_a_*. This started from *c_a_* = 1, with the strength of preferential flow increasing as the value of *c_a_* increased by factors of 10. These simulations were run for 7 soil types on the loam to clay spectrum (Carsel and Parrish, 1988).

For investigating the influence of root architecture on root water uptake, and water lifetime in the rooted zone, the model (1), (8) was parametrised for each of the root systems 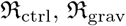, and 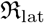. Simulations were obtained from each parametrisation, with the strength of induced preferential flow kept constant at 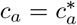 in all three. At the lower boundary Γ_3_, the value of *K_s_* was increased by a factor of 10 to impose a condition of enhanced free drainage. This modelled the situation of having a layer of more hydraulically conductive material at the lower boundary, which increases the rate of drainage, and creates an effect similar to that of run off along bedrock.

### Computations

The conformal finite-element method, with an implicit Euler discretisation in time, was used to obtain numerical pressure head solutions to our model (1), (8) and the benchmark model. The finite-element mesh consisted of a disjoint union of tetrahedra, where the maximum possible circumradius of a tetrahedron was 0.059m, and the minimum was 0.029m. A time step of 0.01d was used in the implicit Euler scheme and the linearisation of the water retention and hydraulic conductivity functions was achieved using an L-method (List and Radu, 2016). This finite-element scheme was implemented using the FEniCS library (Alnæs et al., 2015). The algorithms used to construct functions *ψ*, *H*, and *RLD* were carried out in Python 3 using the NumPy and SciPy libraries (Harris et al., 2020). Root density profiles and simulations of water content evolution were visualised using Paraview (Ahrens et al., 2005), and the plots in Figures 2 (a) and 4, showing the architectures of simulated root systems, were generated using MATLAB 2020a. The Bayesian optimisation algorithm used to find the optimal facilitation constant 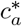, and hence calibrate model (1), (8), was implemented using the scikit-optimise library in Python 3 (Head et al., 2018).

## Results

### Model Calibration

The model (1), (8) was parametrised for the root system 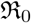 (Figure 2 (a)), and *H* itself was parametrised with a facilitation constant *c_a_*. For a given *c_a_* value, the pressure head profile *h_C_a__* was the numerical solution of the corresponding parametrisation of model (1), (8). A number of Bayesian optimisation schemes with low total iteration numbers were first run to search the interval [1, 350000] for a *c_a_* value that minimised the cost function *u*. Through this method, a trough in the value of *u*(*c_a_*) was identified within the interval [268000, 272000]. Since it is likely that the cost function *u* has a global minimiser (Mair et al., 2022), a further Bayesian optimisation scheme of 15 iterations was run on the interval [268000, 272000]. This scheme identified a minimiser of 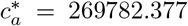, where 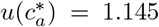. Since the final iterations of the scheme were clustered at points within a small neighbourhood of 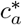, all yielding similarly low cost function values, the scheme was deemed to have sufficiently converged and no further iterations were performed. The close agreement between profiles 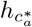 from the model (1), (8), calibrated with 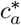, and profiles from the benchmark model is shown by Figures 2 (b) and (c). This indicates that, for Maize root systems, our procedure is indeed capable of calibrating model (1), (8) against experimental data.

### The impact of preferential flow on root water uptake and water loss from the rooted zone

All scenarios considered resulted in zero run off losses. Total water loss from the rooted zone was therefore taken to be the sum of total evaporation and deep percolation. When considering the reference soil (silt loam), it was found that increasing the value of the facilitation constant from *c_a_* = 1 caused total cumulative evaporation losses to decrease (Figure 3 (c)). By contrast, total cumulative deep percolation losses were found to increase as the value of the facilitation constant was increased (Figure 3 (d)). This meant that, total cumulative water loss decreased as the value of the facilitation constant increased from *c_a_* = 1 to a critical value of *c_a_* ≈ 10^6^ (Figures 3 (a)). However, for increases in *c_a_* beyond this critical value, the increase in deep percolation losses, exceeded the decrease in evaporation losses. This resulted in a net increase in water loss from the rooted zone (Figure 3 (a)). Similarly, cumulative root water uptake was found to increase as the facilitation constant was increased from *c_a_* = 1, but then decrease once *c_a_* was increased past some critical value (Figure 3 (b)). The same pattern was observed across all soil types considered, but with different critical *c_a_* values for minimising water loss from the rooted zone, or maximising uptake (Figure 3 (e) and (f)). Changing the strength of the preferential flow induced by the root system had the greatest impact on water loss in loam and sandy clay loam (Figure 3 (e)). Out of the soil types considered, these have the highest values for the saturated hydraulic conductivity *K_s_* and water retention parameter *α*_vg_ (Carsel and Parrish, 1988). On the other hand, for root water uptake, the strength of preferential flow had the largest effect in clay loam, silt and silt loam (Figure 3 (f)). These soil types have *K_s_* and *α*_vg_ values that are closest to the average over all seven types considered. For total water loss and root water uptake, the observed effect of changing the strength of root-induced preferential flow was smallest within clay, and silty clay loam (Figure 3 (e), (f)). These soil types have the lowest values for *K_s_* and *α*_vg_ out of the seven considered.

**Fig. 3:**
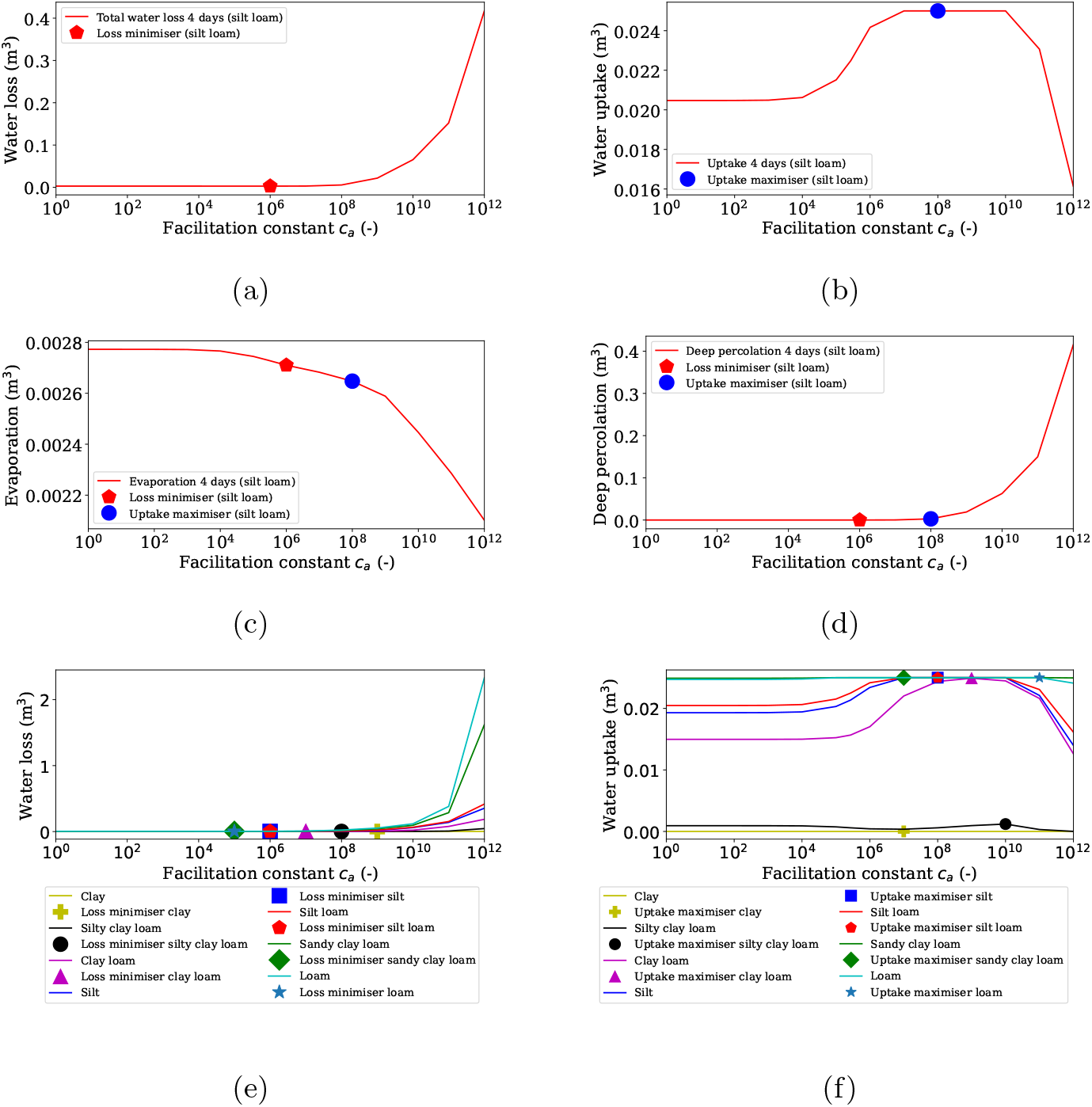
The influence on water lifetime in the rooted zone from increasing and decreasing the strength of preferential flow induced by a root system. (a) Total water losses at increasing facilitation constant values *c_a_* in silt loam soil. (b) Cumulative root water uptake at increasing *c_a_* values in silt loam soil. (c) Water losses from evaporation at increasing *c_a_* values in silt loam soil. (d) Water losses from deep percolation at increasing *c_a_* values in silt loam soil. (e) Total water losses at increasing *c_a_* values for all soil types. (f) Cumulative root water uptake at increasing *c_a_* values for all soil types.

**Fig. 4:**
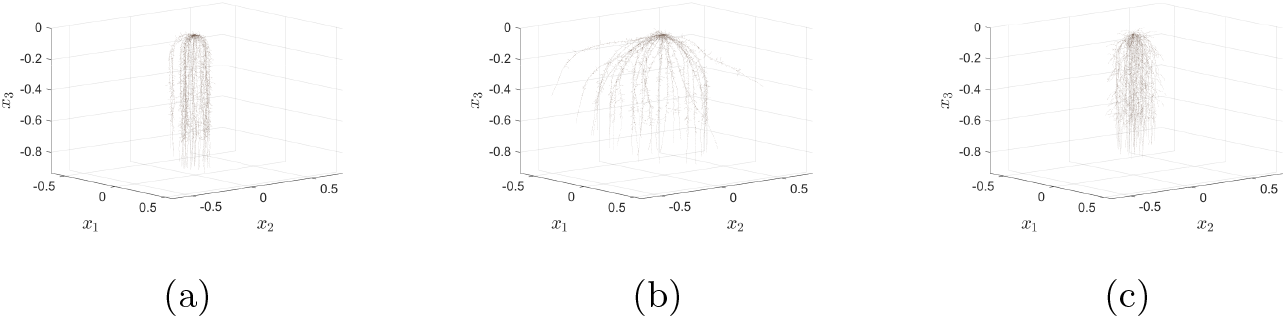
Root systems used for testing the impact of root architectural traits on water loss from vegetated soil. (a) Control root system 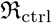. (b) Root system where gravitropism of roots is decreased 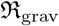. (c) Root system where length of lateral roots is increased 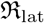. The volumes that these root systems occupy are all within a 2% tolerance of each other.

### Impact of root system traits on soil water losses and uptake

Simulations from model (1), (8) showed zero runoff losses for the soils vegetated by the contrasting root systems in Figure 4. Therefore, total water losses were a sum of evaporation and deep percolation. Results showed that the cumulative total water loss from the rooted zone was greatest when the soil was vegetated by the control root system 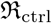, and lowest when vegetated by the root system 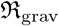, in which the gravitropism of roots had been reduced (Table 1). Over the entire simulation period, the rate of total water loss from the rooted zone (m^3^d^-1^) was lowest when vegetated by 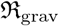, and highest when vegetated by 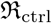 (Figure 5 (a)).

**Fig. 5:**
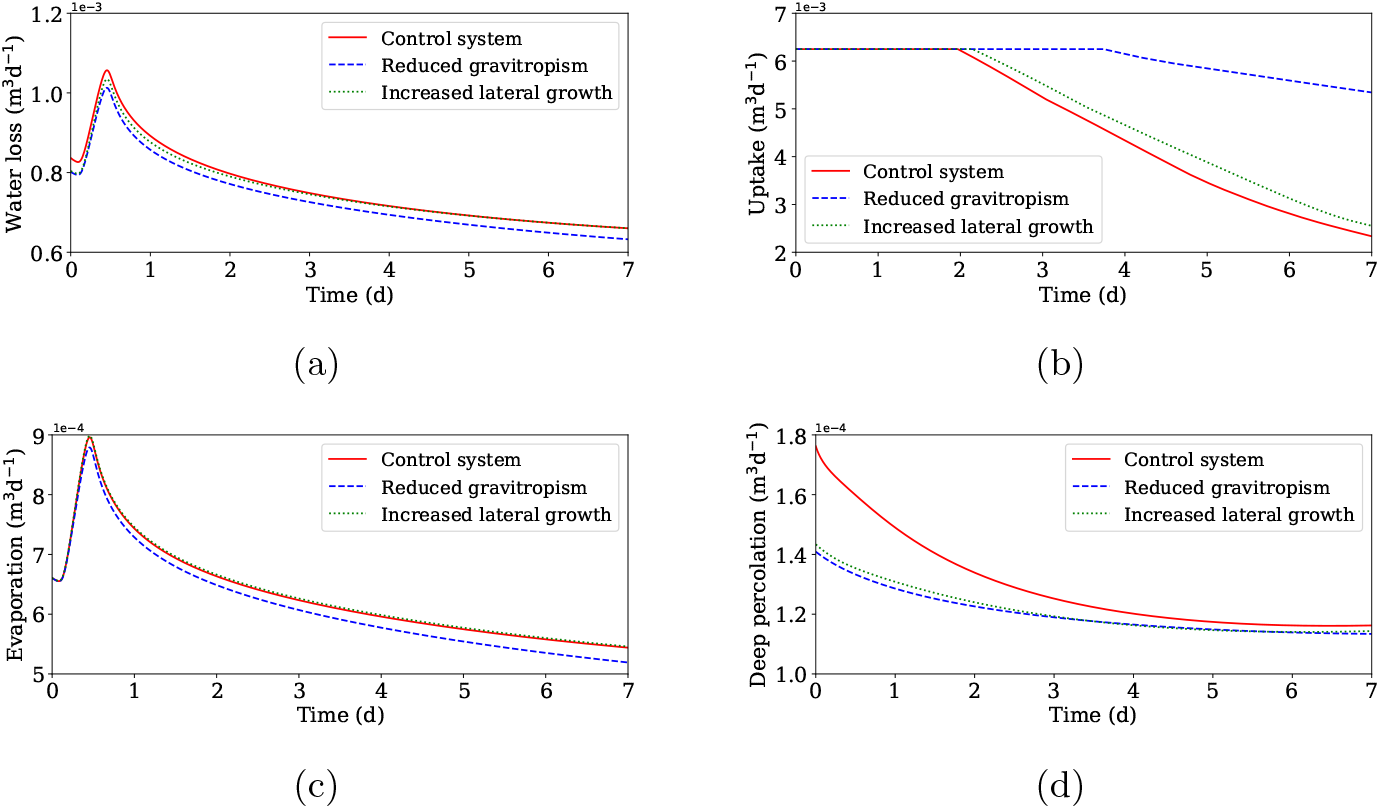
Water losses and uptake rate from the rooted zone following precipitation for 3 identically shaped soil domains, each vegetated by one of the root systems shown in Figure 4. (a) Rate of total water loss from the rooted zone over the 7 day period. (b) Rate of total water uptake for each root system over the 7 day period. (c) Rates of water loss from the rooted zone via evaporation over the 7 day period. (d) Rates of water loss from the rooted zone via deep percolation over the 7 day period.

For each of the soils vegetated by the 3 different root systems, approximately 15 – 20% of cumulative water loss was attributed to deep percolation. The losses via deep percolation were greatest within the soil vegetated by 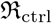. The soils vegetated by 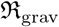, and the system with longer lateral roots 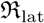, both exhibited lower values for cumulative deep percolation losses (Table 1). In fact, deep percolation rate was highest, at all times of the simulation, within the soil vegetated by 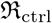. This was particularly noticeable at early simulation times (Figure 5 (d)). To a lesser degree, the rate of deep percolation was higher when the soil was vegetated by 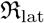, as opposed to 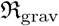. Cumulative deep percolation losses were also marginally greater when the soil was vegetated by 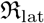, instead of 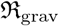 (Table 1).

Regardless of root system, the majority of total soil water loss was always due to evaporation (Table 1). For the entire duration of the simulation, the rate of water loss via evaporation was lowest in the soil that was vegetated by 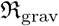 (Figure 5 (a)). However, there was little difference, at any point in the simulation, between the rates of evaporation from the soils vegetated by 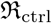, and 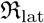 (Figure 5 (a)). In terms of cumulative evaporation, the soil vegetated by 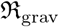 again lost the least. However, the soil vegetated by 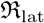 actually lost marginally more water via evaporation than the soil vegetated by 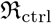 (Table 1).

The water uptake performance of each root system followed an unsurprising trend. The cumulative root water uptake of 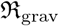 was 29% higher than 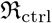, and 23% higher than 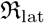. Furthermore 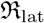 achieved a cumulative water uptake that was 4% higher than 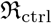 (Table 1). Further insight into these cumulative results is provided by the uptake rates of each root system over the seven day period (Figure 5 (b)). On the first two days, the root systems were all taking up water at the same maximum rate. On day 3, the water uptake rate of 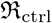 began to decline. This was followed shortly after by the uptake rate of 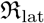, albeit with a shallower rate of decline. It was not until late in day 4 that the rate of uptake of 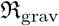 started to decline, and this decline was considerably more shallow than for the other two root systems.

It was observed (Figures 6 and 7) that each of the root systems in Figure 4 induced different water flux patterns within the soil. Root-induced redistribution of soil water was driven by preferential flow, and also the pressure gradients that arose from the removal of water by root uptake. At early stages of the simulation, the dominant effect of root system 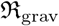, on soil water distribution, was to facilitate increased infiltration from the soil surface into the bulk soil (Figure 6 (d)). This effect was observed across a larger area of the upper soil surface than in the soils vegetated by 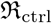 or 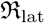 (Figures 7 (a), (d) and (g)). For both 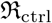 and 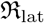, the redistribution of soil water at early stages of the simulation was driven mainly by root water uptake from within the narrower regions of soil that they occupied (Figures 6 (a), (g)).

**Fig. 6:**
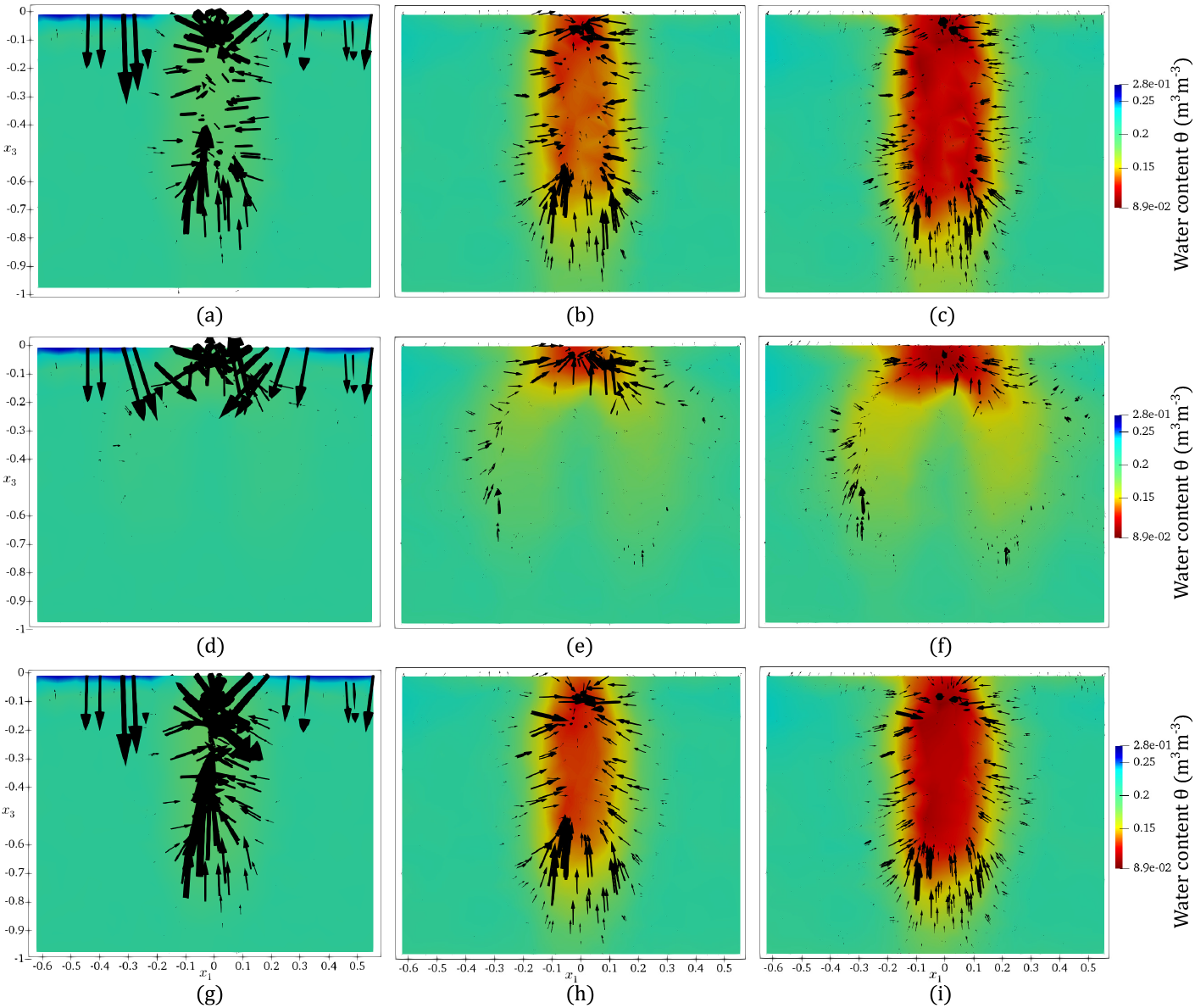
Evolution of post precipitation soil water content in the rooted zone. Three identical silt loam domains, each vegetated with one of the root systems in Figure 4. Plots (a), (d), (g) show water content profiles after 0.5 days for soils vegetated by the control 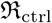, the reduced gravitropism 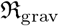, and the increased lateral growth 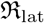 root systems respectively. Plots (b), (e), (h) show water content profiles after 3.75 days for soils vegetated by the root systems 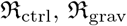, and 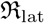 respectively. Plots (c), (f), (i) show water content profiles after 7 days for soils vegetated by the root systems 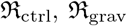, and 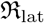 respectively. Arrows indicate the direction and strength of the water flux in different regions of the domain.

**Fig. 7:**
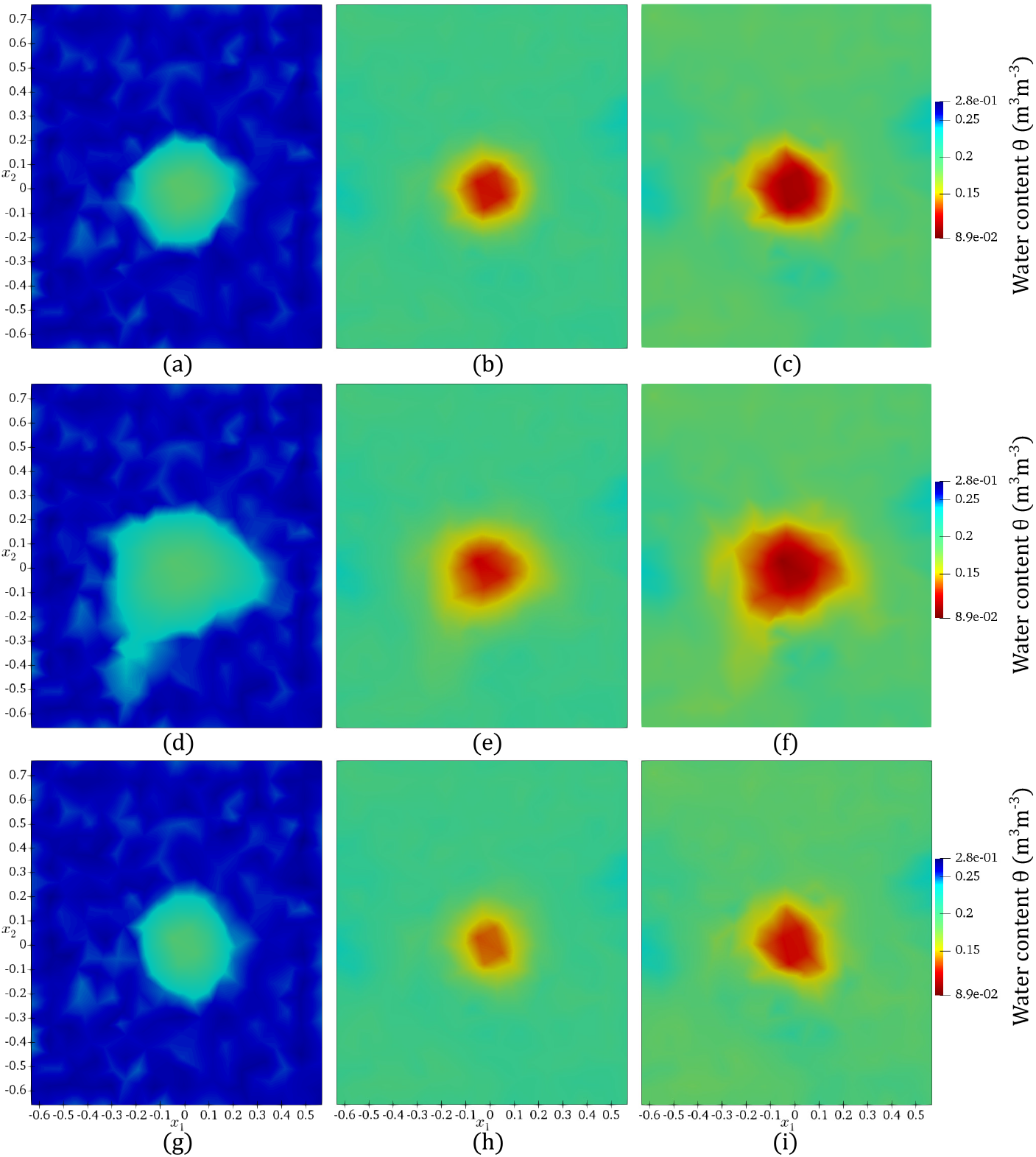
Evolution of post precipitation soil water content in the rooted zone of three identical silt loam domains, each vegetated with one of the root systems in Figure 4. The view looks down on the upper soil surface. Plots (a), (d), (g) show water content profiles after 0.5 days for soils vegetated by the control 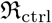, the reduced gravitropism 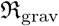, and the increased lateral growth 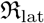 root systems respectively. Plots (b), (e), (h) show water content profiles after 3.75 days for soils vegetated by the root systems 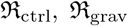, and 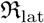 respectively. Plots (c), (f), (i) show water content profiles after 7 days for soils vegetated by the root systems 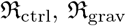, and 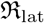 respectively.

At all simulation times, the influence of 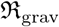, on the orientation and magnitude of soil water flux, spanned a greater lateral area than both 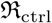 and 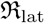 (Figures 6, 7). This meant that, during later stages of the simulation, 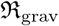 had facilitated the infiltration into bulk soil of a greater proportion of surface water than either of the other two root systems (Figure 7). Consequently, in the period from the end of the precipitation event to the end of the simulation, the water content of the bulk soil around 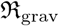 was higher than in the bulk soil around the other two root systems (Figure 6).

## Discussion

### The trade-off between evaporation losses and losses by deep percolation

The environmental conditions experienced by many farmed crops, lead to total water losses from the vegetated soil being dominated by evaporation, with a smaller proportion attributed to deep percolation (Paruelo and Sala, 1995;Schwinning and Sala, 2004). An increase in infiltration from the soil surface into the bulk soil will therefore reduce the amount of surface water vulnerable to evaporation, without equivalently increasing deep percolation losses. This means that more water is available to the root system, and for longer, which delays the emergence of water deficits, and allows the plant to maintain a healthy rate of water uptake during periods of no precipitation.

In our model (1), (8), the strength of preferential flow induced by a root system can be increased by increasing the value of the facilitation constant *c_a_*. Simulations showed that the values of facilitation constant that minimised total water loss, and maximised root water uptake, were orders of magnitude higher than the facilitation constant 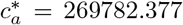, which was estimated from experimental data for Maize (Figures 3). This indicates that Maize root systems may not increase infiltration into the bulk soil to an extent that is sufficient to minimise water loss. However, results also indicated that if the strength of root-induced preferential flow was increased too far, then the consequent increase in deep percolation losses would exceed the reduction in evaporation losses, and cause a net increase in total water loss (Figure 3). Similar results were obtained for all soil types considered, and showed that root systems in soils with higher hydraulic conductivity did not need to induce as strong a preferential flow, in order to minimise water loss and maximise uptake (Figure 3 (e), (f)).

Considering experimental research to date, there remains some doubt as to which root traits promote preferential flow through soil. However, potential candidates include root hair growth, the exudation of mucilage by root tissue, and the activity of the rhizosphere-microbiome, which have all been shown to affect the hydraulic characteristics of vegetated soil (Hallett et al., 2022; Carminati et al., 2010; Choudhury et al., 2018). Through genotyping, it is possible to produce phenotypes that vary in their expression of these traits (Hochholdinger and Tuberosa, 2009; Ahmad et al., 2011; Bilyera et al., 2021). This suggests that selecting specific phenotypes, which induce preferential flow to an extent that is optimal for extending the lifetime of water in the soil type they inhabit, may provide a new strategy for breeding drought resistant crops.

### The effect of root system architecture on water lifetime in vegetated soil

Following precipitation, root-oriented preferential flow influences patterns of infiltration into soil (Noguchi et al., 1997). Because of this, root systems with differing architectures are likely to induce different distributions of water throughout the soil. The presence of roots near the surface of soil has been observed in experiments to increase infiltration rates into bulk soil (Song et al., 2017; Leung et al., 2018). This increased infiltration can be attributed initially to roots near the soil surface inducing preferential downward flow, then subsequently to roots removing water from the soil via uptake, and steepening the pressure head gradient between upper and lower soil layers. The mass of the reduced gravitropism root system 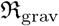 is distributed over a larger lateral area than the other systems (Figure 4). This was particularly true at shallower soil depths. Therefore, by the mechanisms mentioned above, 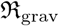 increased infiltration, from surface soil to bulk, across a larger lateral area of the domain (Figure 7). For this reason, less soil water was left at the surface, in comparison to the other root systems, hence less water was lost to evaporation. Water losses via deep percolation were lowest in the soil vegetated by 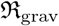, and highest, by a considerable margin, in the soil vegetated by the control root system 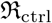 (Table 1). Reduced gravitropism and longer first-order laterals, mean a smaller proportion of total root length is oriented vertically downward, than in a system with typical root architecture like 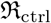 (Figure 4). Consequently these root systems divert more soil water transport into lateral directions than 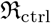, and less water will pass the lower boundary of the rooted zone. Secondly, if roots take up water, then that water is not lost via deep percolation. Moreover, the removal of water via uptake will steepen pressure head gradients within the soil and drive more capillary rise. Compared to 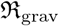, root water uptake by 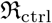 and 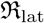 was concentrated within a narrower region of the soil (Figures 6 and 7). As a result, capillary rise was induced within a narrower region of the soil (Figure 6), and more water was lost from the soil via deep percolation.

Since the soil with the root system 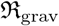 lost the least water to evaporation and deep percolation (Figure 5, Table 1), it is unsurprising that this root system maintained the highest uptake rate at all simulation times. We do however propose one more contributing factor toward this result. Since the roots of 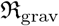 span a larger volume of the soil, uptake is not limited to a narrow soil region, like it is for 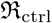 and 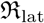 (Figures 6, 7). This means that the water available for uptake is depleted less quickly by 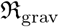 thus allowing it to maintain a higher uptake rate. All these findings lead us to the conclusion that, in scenarios involving intermittent precipitation events followed by periods of drought, Maize root systems with reduced gravitropism (Figure 4 (b)) are most adept at extending water lifetime in the rooted zone and maximising uptake efficiency.

### Root system ideotypes for drought resistance

Numerous morphological and physiological features of plant roots are involved in the acquisition of water and nutrients from soil. The length and density of root hairs is critical to the uptake of immobile nutrients such as phosphorous and potassium (Jungk, 2001). For accessing mobile resources such as nitrogen and water, structural traits like root growth angle have been shown to play a crucial role (Uga et al., 2013). In terms of physiological adaptations, experimental investigations have revealed that some plants form root cortical aerenchyma as a response to the effects of drought conditions (Zhu et al., 2010) and soil water logging (Yamauchi et al., 2018). Furthermore, the exudation of mucilage by plant roots is known to influence the composition and activity of the root microbiome (Badri and Vivanco, 2009). This affects the hydraulic characteristics of the rhizosphere (de la Fuente Cantó et al., 2020), and the radial hydraulic conductivity of the root tissue, which in turn influences the capacity of the root to absorb water and nutrients (Pierret, 2022).

There is some disagreement as to exactly what a root ideotype for water use efficiency and drought resistance should look like (Tardieu, 2012; Tron et al., 2015; Pierret, 2022). The influential study of Lynch (2013) concluded that drought resistant root systems must be “steep, cheap and deep”(SCD) i.e. have large diameter primary roots that grow deep with fewer, but longer, laterals (Nepstad et al., 1994; Schenk and Jackson, 2002; Lynch and Wojciechowski, 2015) and this was later supported by the experimental results of Uga et al. (2013). The justification for this comes from the premise that such an architecture allows access to water stored at greater depths, with the presence of some root mass near the soil surface meaning uptake at shallower depths is not overly compromised (Lynch, 2013). However, when considering an initial condition of higher water content near the soil surface, the simulations of Leitner et al. (2014) found that an SCD maize root system achieved a lower cumulative water uptake than one with a standard structure. Lynch (2013) also concede that in semi-arid climates, with intermittent periods of drought punctuated by regular rainfall events, the advantage of an SCD root system structure is less obvious. Furthermore, recent work by Clément et al. (2022) has shown that the axial hydraulic conductivity of root tissue decreases with depth, and that SCD root systems may therefore be unable to effectively utilize all the deeply stored water that their architectures provide them access to.

Since soil water content is very dynamic, we propose a departure from the approach to identifying root system ideotypes for drought resistance that focuses solely on the spatial distribution of roots, and the capacity to access static stores of soil water. Instead, we emphasise the capacity of root systems to induce preferential flow patterns, as a result of their architecture and physiological activity, which increase the lifetime of water in the rooted soil by decreasing losses via evaporation and deep percolation. This incorporates traits that influence the characteristics of the surrounding soil into the concept of a root system ideotype for drought resistance. Such a perspective is an example of the concept of an extended phenotype for drought resistance (de la Fuente Cantó et al., 2020), and for plants that feed on water accumulated at great depths, it is still likely to identify the SCD root system as optimal (Lynch,2013). However, if crops are rain fed or irrigated by sprinklers, and experience periods of intermittent drought, then root systems with reduced gravitropism may indeed make the most efficient use of water. Further experimental research is required to validate the conclusions of this work, and provide more evidence that root-oriented preferential flow is a crucial factor to consider, when identifying crop ideotypes for drought resistance.

## Acknowledgements

Andrew Mair was supported by The Maxwell Institute Graduate School in Analysis and its Applications, a Centre for Doctoral Training funded by the UK Engineering and Physical Sciences Research Council (grant EP/L016508/01), the Scottish Funding Council, Heriot-Watt University and the University of Edinburgh. Lionel Dupuy was supported by the Spanish Ministry of Science and Innovation (MICINN) under the project MICROCROWD (PID2020-112950RR-I00).

## Author Contributions

AM derived the mathematical model (1), (8), coded and carried out the simulation method, sourced the paper by Feki et al. (2018) from which experimental results were used, designed, coded and carried out the optimisation method to calibrate the mathematical model against experimental data, coded the scripts to run the simulations of the scenarios investigated, performed the programming/formatting required to display results visually, created the concept figure (Figure 1), wrote the original draft of the manuscript and all subsequent versions (after review by both supervisors).

LD provided supervision, gave advice on model derivation and boundary conditions, put forward the specific infiltration scenarios we should investigate through simulation, reviewed and edited drafts of the paper, and recommended references.

MP provided supervision, assisted in model derivation and the choice of numerical method, reviewed and edited drafts of the paper, and recommended references.

All authors gave final approval for publication and agree to be held accountable for the work performed therein.

## Data Availability

Data and code used to generate the results and figures from this submission can be accessed via a Dropbox folder that will be set up by the Open Research Data team at Heriot-Watt University. Reviewers and editors should contact Linda Kerr, or the Open Research Data team, at Heriot-Watt University and they will be given access. Within the folder is a “readme file”. The text files “fenics_setup” and “operating_procedure” should be used as a guide to recreate the results of this submission. The full link to a DOI containing the data and code will be provided if the manuscript is published in a scientific journal.

## Notes

### Competing Interest Statement

The authors have declared no competing interest.

