## Supplementary Material for "Can root systems redistribute soil water to mitigate the effects of drought?"

### Model

#### Root water uptake and transpiration

The function for root water uptake in the main text is given as

$$S(x,t) = \alpha_{\mathrm{F}}(h) NRLD(x) A_{\mathcal{T}} \mathcal{T}_{\mathrm{p}} \frac{1}{\max\left(\omega(t), \omega_c\right)}. \tag{1}$$

The dimensionless water stress response function $\alpha_{\mathrm{F}}$, has the following form (Feddes, 1982):

$$\alpha_{\mathrm{F}}(h) = \begin{cases} 0 & h \geq h_1,\ h \leq h_4, \\ \frac{h-h_1}{h_2-h_1} & h_2 \leq h < h_1, \\ 1 & h_3 < h \leq h_2, \\ \frac{h-h_4}{h_3-h_4} & h_4 < h \leq h_3. \end{cases} \tag{2}$$

In this formulation, $h_1 < 0$ is the pressure head associated with plant anaerobiosis occurring at high levels of soil water content, and $h_4 < 0$ is the pressure head corresponding to the wilting point of the plant at low soil water values. The values of $h_2$ and $h_3$ give the pressure head interval within which root water uptake is unstressed, and maximum uptake rate is maintained. Outwith the interval $(h_3, h_2]$ the water stress on the plant increases and root water uptake decreases. This is reflected by the water stress response function $\alpha_{\mathrm{F}}$ which decreases linearly to 0 as $h$ goes to $h_1$ or $h_4$. The lower bound $h_3$, of the pressure head interval within which root water uptake assumes its maximum rate, is taken to be a piecewise linear function which depends upon potential transpiration $\mathcal{T}_{\mathrm{p}}$ (Cai et al., 2018):

$$h_3(\mathcal{T}_{\mathrm{p}}) = \begin{cases} h_{3h} & \mathcal{T}_{\mathrm{p}} \geq \mathcal{T}_{\mathrm{3h}}, \\ h_{3h} + \frac{(h_{3l}-h_{3h})(\mathcal{T}_{\mathrm{3h}}-\mathcal{T}_{\mathrm{p}})}{\mathcal{T}_{\mathrm{3h}}-\mathcal{T}_{\mathrm{3l}}} & \mathcal{T}_{\mathrm{3l}} < \mathcal{T}_{\mathrm{p}} < \mathcal{T}_{\mathrm{3h}}, \\ h_{3l} & \mathcal{T}_{\mathrm{p}} \leq \mathcal{T}_{\mathrm{3l}}. \end{cases} \tag{3}$$

The value of $h_3$ decreases linearly to $h_{3l}$ as transpiration decreases toward a low rate $\mathcal{T}_{\mathrm{3l}}$ ($\mathrm{LT}^{-1}$), but increases linearly to $h_{3h}$ as the transpiration rate increases to a high value $\mathcal{T}_{\mathrm{3h}}$. This relationship, modelled by (3), reflects the fact that the difference in pressure between shoot and root is greater under higher transpiration rates (Cai et al., 2018).

Potential transpiration is given by the formula $\mathcal{T}_{\mathrm{p}} = \mathrm{K_{cb}ET_o}$. Here $\mathrm{ET_o}$ is the reference evapotranspiration ($\mathrm{LT}^{-1}$), and total evapotranspiration ($\mathrm{LT}^{-1}$) is calculated as $\mathrm{ET_c} = (\mathrm{K_{cb}} + \mathrm{K_e})\mathrm{ET_o}$ (Allen et al., 1998). The contribution to total evapotranspiration coming from crop transpiration is then determined by the value of the basal crop coefficient $\mathrm{K_{cb}}$, and $\mathrm{K_e}$ dictates the amount of evaporation from the soil surface. Typical values of $\mathrm{K_{cb}}$ for different plant species and seasons can be found in Table 17 Chapter 7 of Allen et al. (1998). The expression for $\mathrm{K_e}$ is given in a following section.

The normalised root length density $NRLD$ incorporates the dependence of root water uptake on the distribution of roots. This function is formulated so that it integrates to unity over the soil domain, and therefore acts as a weighting function that determines the extent to which a region of soil contributes to uptake and transpiration. The soil surface area associated with transpiration is denoted as $A_{\mathcal{T}}$ ($L^2$), and the dimensionless water stress index $\omega(t) = \int_\Omega \alpha_F(h(x,t))NRLD(x)dx$, is the ratio between total uncompensated transpiration $\mathcal{T}_a(t) = \mathcal{T}_p \int_\Omega \alpha_F(h(x,t))NRLD(x)dx$ and potential transpiration $\mathcal{T}_p$ (Simunek and Hopmans, 2009). The term $(\max(\omega, \omega_c))^{-1}$ is what incorporates compensated uptake into $S$, and with actual compensated transpiration denoted as $\mathcal{T}_{ac} = A_{\mathcal{T}}^{-1} \int_\Omega S(x,t)dx$, its action is described as follows:

$$\begin{aligned}
\omega = 1 &\implies S(x,t) = \alpha_F(h)NRLD(x)A_{\mathcal{T}}\mathcal{T}_p &&\implies \mathcal{T}_a = \mathcal{T}_{ac} = \mathcal{T}_p,\\
\omega_c \le \omega < 1 &\implies S(x,t) = \frac{\alpha_F(h)NRLD(x)A_{\mathcal{T}}\mathcal{T}_p}{\int_\Omega \alpha_F(h(x,t))NRLD(x)dx} &&\implies \mathcal{T}_a < \mathcal{T}_{ac} = \mathcal{T}_p,\\
\omega < \omega_c &\implies S(x,t) = \frac{\alpha_F(h)NRLD(x)A_{\mathcal{T}}\mathcal{T}_p}{\omega_c} &&\implies \mathcal{T}_a < \mathcal{T}_{ac} < \mathcal{T}_p.
\end{aligned} \tag{4}$$

It can be seen from (4) that if $\omega = 1$, then the plant is not experiencing any water stress so the actual transpiration rate meets the potential transpiration rate without the need for any compensation. If $\omega_c \le \omega < 1$, then the plant is experiencing some water stress, but the actual compensated transpiration rate still meets the potential transpiration rate due to the root system adapting and increasing root water uptake. This increase is proportional to $\alpha_F$ and hence greater in regions of rooted soil that are wetter. If $\omega < \omega_c$, then compensation occurs, but the global level of water stress experienced by the plant is too large for this to bring the actual compensated transpiration rate in line with the potential transpiration rate (Simunek and Hopmans, 2009; Cai et al., 2018).

#### Boundary functions

The term $K_e$ that governs evaporation from the upper soil surface is defined as a function of pressure head $h$ (Allen et al., 1998):

$$K_e(h) = \min[K_r(h)(K_{cmax} - K_{cb}), f_{ew}K_{cmax}]. \tag{5}$$

Here $K_{cmax}$ is the maximum value that the crop coefficient $K_c = K_{cb} + K_e$ can take after precipitation, and $f_{ew} = \min(1 - f_c, f_w)$ is the fraction of the soil surface that is wetted and exposed. The constants $f_c$, $f_w \in [0.1, 1]$ give the average fractions of the upper soil surface that are covered by vegetation and wetted by precipitation respectively. The function $K_r$ is the evaporation reduction coefficient and governs the relationship between the amount of evaporation taking place and the water content at the soil surface:

$$K_r(h) = \begin{cases} 0 & \theta(h) \le 0.5\theta_{WP} \\ \frac{\theta(h) - 0.5\theta_{WP}}{\theta_{FC} - 0.5\theta_{WP}} & 0.5\theta_{WP} < \theta(h) < \theta_{FC} \\ 1 & \theta(h) \ge \theta_{FC}, \end{cases} \tag{6}$$

where $\theta_{\mathrm{WP}}$ and $\theta_{\mathrm{FC}}$ are the water contents at wilting point and field capacity respectively (Allen et al., 1998). The precipitation condition used for calibrating the model (Figure 1) imposed a rainfall event on each day of the 2 day period that was simulated:

$$\mathrm{P}_0(t) = 0.0052\left[\left(1+\exp\left(-2\left(1-\frac{(t-0.3)^2}{0.125^2}+2^{-\frac{1}{2}}\right)\right)\right)^{-1} + \left(1+\exp\left(-2\left(1-\frac{(t-1.3)^2}{0.125^2}+2^{-\frac{1}{2}}\right)\right)\right)^{-1}\right], \qquad t \geq 0. \tag{7}$$

The precipitation condition for investigating the impact of preferential flow and root architectural traits on water loss from the rooted soil (Figure 2), prescribed a rainfall only on day 1 of the time period simulated:

$$\mathrm{P}(t) = 0.0052\left(1+\exp\left(-2\left(1-\frac{(t-0.3)^2}{0.125^2}+2^{-\frac{1}{2}}\right)\right)\right)^{-1}, \qquad t \geq 0. \tag{8}$$

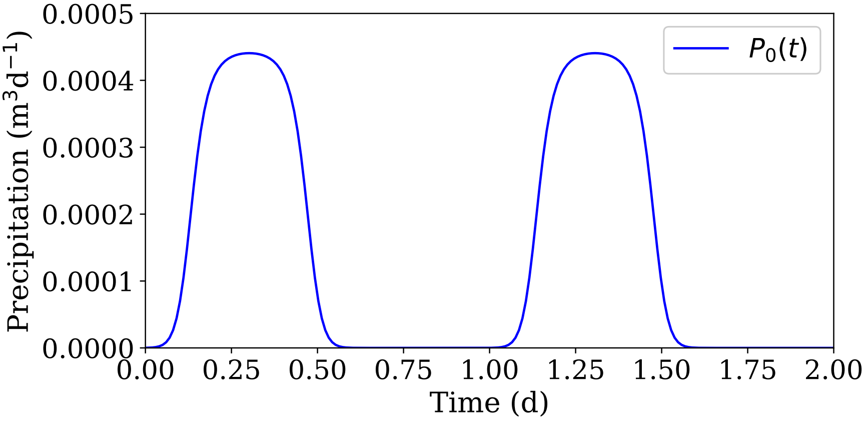


**Fig. 1**: The precipitation event considered in the scenario used for calibration.

#### Benchmark model

The experimental data of Feki et al. (2018) provides a value $K_s^*$ for the saturated hydraulic conductivity of soil vegetated by Maize. This allows the formulation of a depth dependent saturated hydraulic conductivity $\hat{K}_s$. Here $\hat{K}_s = K_s^*$ in the vegetated section of the soil domain above rooting depth, and $\hat{K}_s = K_s$ in the fallow soil below. Like in (Mair et al., 2022), this then

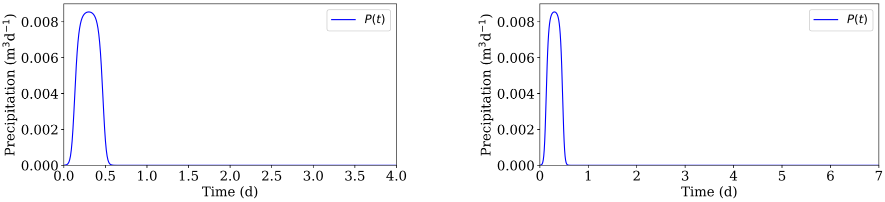


(a) (b)

**Fig. 2**: Precipitation events used in the investigations into the influence of preferential flow, and root architectural traits, on water lifetime in the rooted zone and uptake efficiency. (a) Precipitation event in the investigation into the influence of preferential flow (b) Precipitation event in the investigation that incorporates the influence of root architectural traits.

results in a benchmark model for the influence of a Maize root system on water infiltration through soil:

$$\begin{aligned} \partial_t\theta(h) + \boldsymbol{\nabla}\cdot\hat{\mathbf{q}}(x) &= -S(x,t), && \text{in } \Omega\times(0,T], \\ \hat{\mathbf{q}}\cdot\mathbf{n} &= \hat{q}_{\mathrm{up}} && x\in\Gamma_1,\ t\in(0,T], \\ \hat{\mathbf{q}}\cdot\mathbf{n} &= 0 && x\in\Gamma_2,\ t\in(0,T], \\ \hat{\mathbf{q}}\cdot\mathbf{n} &= \hat{q}_{\mathrm{low}} && x\in\Gamma_3,\ t\in(0,T], \\ h(x,0) &= h_0(x) && x\in\Omega, \end{aligned} \tag{9}$$

with

$$\hat{\mathbf{q}}(x) = -\hat{K}_s\kappa(h)\boldsymbol{\nabla}(h+x_3), \tag{10}$$

and

$$\hat{q}_{\mathrm{low}} = K_s\kappa(h). \tag{11}$$

Here the upper boundary condition is $\hat{q}_{\mathrm{up}} = \tilde{q}_{\mathrm{up}}$, and full expressions for the functions $\tilde{q}_{\mathrm{up}}$, $h_0$, $\theta$, $\kappa$ and $S$ are given in the main text.

#### Cost function for calibration

A cost function $u$ was minimised to identify the facilitation constant $c_a^*$ with which to parametrise the model from the main text, so that it best reflected empirical data of Feki et al. (2018). With the numerical pressure head solution from the benchmark model given as $h_b$, the corresponding laterally averaged pressure head profile is given by:

$$\overline{h}_{\mathrm{b}}(x_3,t) = \frac{1}{|\mathcal{A}_{\mathrm{lat}}|}\int_{\mathcal{A}_{\mathrm{lat}}} h_b(x_1,x_2,x_3,t)dx_1dx_2, \tag{12}$$

Where $\mathcal{A}_{\mathrm{lat}} \subset \mathbb{R}^2$ is the surface defined by the maximum and minimum possible values of $x_1$ and $x_2$ for any position $x\in\Omega$. In evaluating $u(c_a)$ for a given $c_a$ value, the volumetric density function $\psi$ and flow anisotropy matrix $H(c_a)$ were first constructed for the root system $\mathfrak{R}_0$ to obtain a parametrisation of the model from the main text. Laterally averaged pressure head solutions to this parametrisation, denoted by $\overline{h}_{c_a}$, were then obtained. With discrete time points $t_1 = 0.5$d, $t_2 = 1$d $t_3 = 1.5$d, $t_4 = T = 2$d, and a number $N_D$ of equally spaced soil depths $\{x_3^1, ..., x_3^{N_D}\}$, $u(c_a)$ was given as the sum of the squared euclidean distance, at each discrete time and depth

$$u(c_a) = \sum_{j=1}^{4} \sum_{i=1}^{N_D} \left[ \overline{h}_c(x_3^i, t_j) - \overline{h}_b(x_3^i, t_j) \right]^2. \tag{13}$$

| Root system | $\mathfrak{R}_0$ | $\mathfrak{R}_{\text{pf}}$ | $\mathfrak{R}_{\text{ctrl}}$ | $\mathfrak{R}_{\text{grav}}$ | $\mathfrak{R}_{\text{lat}}$ |
|---|---|---|---|---|---|
| Seed location | (0,0,0) | (0,0,0) | (0,0,0) | (0,0,0) | (0,0,0) |
| Days growth | 90 | 90 | 90 | 90 | 90 |
| Tropism parameter 1 (mean): primary roots | 1.5 | 1.5 | 1.5 | 0.75 | 1.5 |
| Tropism parameter 1 (mean): 1st order laterals | 1 | 1 | 1 | 0.5 | 1 |
| Tropism parameter 1 (mean): 2nd order laterals | 2 | 2 | 2 | 1 | 2 |
| Tropism parameter 2 (mean): primary roots | 0.16 | 0.16 | 0.16 | 0.08 | 0.16 |
| Tropism parameter 2 (mean): 1st order laterals | 0.31 | 0.31 | 0.31 | 0.155 | 0.31 |
| Tropism parameter 2 (mean): 2nd order laterals | 0.31 | 0.31 | 0.31 | 0.155 | 0.31 |
| Max length (mean): primary roots (m) | 0.897 | 0.897 | 0.897 | 0.897 | 0.808 |
| Max length (mean): 1st order laterals (m) | 0.006 | 0.006 | 0.006 | 0.006 | 0.096 |

| | | | | | |
|---|---|---|---|---|---|
| Inter-lateral distance (mean): primary roots (m) | 0.006 | 0.006 | 0.006 | 0.006 | 0.036 |
| Inter-lateral distance (mean): 1st order laterals (m) | 0.004 | 0.004 | 0.004 | 0.004 | 0.008 |
| Min $x_1$ (m) | $-0.129$ | $-0.149$ | $-0.211$ | $-0.460$ | $-0.185$ |
| Max $x_1$ (m) | 0.168 | 0.157 | 0.161 | 0.566 | 0.231 |
| Min $x_2$ (m) | $-0.133$ | $-0.176$ | $-0.21$ | $-0.657$ | $-0.23$ |
| Max $x_2$ (m) | 0.162 | 0.149 | 0.145 | 0.761 | 0.212 |
| Root depth $D_{\mathfrak{R}}$ (m) | 0.917 | 0.925 | 0.912 | 0.855 | 0.865 |
| Volume ($m^3$) | $3.11\times10^{-5}$ | $3.04\times10^{-5}$ | $3.09\times10^{-5}$ | $3.08\times10^{-5}$ | $3.05\times10^{-5}$ |

**Table 1**: CRootBox simulation settings and data. The settings used to simulate each studied root system from CRootBox (Schnepf et al., 2018), and the dimensions of each generated root system. The root system used for calibrating the model is $\mathfrak{R}_0$. The system used to investigate the influence of preferential flow on RWU, and moisture loss from the rooted zone, is $\mathfrak{R}_{\text{pf}}$. The systems $\mathfrak{R}_{\text{ctrl}}$, $\mathfrak{R}_{\text{grav}}$, and $\mathfrak{R}_{\text{lat}}$ are those which were used to investigate the influence of root architectural traits on RWU, and moisture loss from the rooted zone.

| Scenario | Calibration | Preferential Flow | Architectural Traits | Reference |
|---|---|---|---|---|
| Root system(s) | $\mathfrak{R}_0$ | $\mathfrak{R}_{\text{pf}}$ | $\mathfrak{R}_{\text{ctrl}}$, $\mathfrak{R}_{\text{grav}}$, $\mathfrak{R}_{\text{lat}}$ | N/A |
| Upper /lower surface | $\mathcal{A}_0 = [-0.129, 0.168] \times[-0.133, 0.162]$ | $\mathcal{A}_{\text{gbl}} = [-0.632, 0.566] \times[-0.657, 0.761]$ | $\mathcal{A}_{\text{gbl}} = [-0.632, 0.566] \times[-0.657, 0.761]$ | N/A |
| Domain depth (m) | 1.835 | 0.983 | 0.983 | N/A |

| | | | | |
|---|---|---|---|---|
| Final time $T$ (days) | 2 | 4 | 7 | N/A |
| Soil type | Silt loam | Multiple | Silt loam | N/A |
| $K_s$ ($\mathrm{md}^{-1}$) | 0.108 | Multiple | 0.108 | Carsel and Parrish (1988) |
| $\theta_r$ ($\mathrm{m}^3\mathrm{m}^{-3}$) | 0.067 | Multiple | 0.067 | Carsel and Parrish (1988) |
| $\theta_s$ ($\mathrm{m}^3\mathrm{m}^{-3}$) | 0.45 | Multiple | 0.45 | Carsel and Parrish (1988) |
| $\alpha_{\mathrm{vg}}$ ($\mathrm{m}^{-1}$) | 2 | Multiple | 2 | Carsel and Parrish (1988) |
| $n$ (-) | 1.41 | Multiple | 1.41 | Carsel and Parrish (1988) |
| $h_1$ (m) | -0.15 | -0.15 | -0.15 | Wesseling (1991) |
| $h_2$ (m) | -0.3 | -0.3 | -0.3 | Wesseling (1991) |
| $h_{3\mathrm{h}}$ (m) | -3.25 | -3.25 | -3.25 | Wesseling (1991) |
| $h_{3\mathrm{l}}$ (m) | -6 | -6 | -6 | Wesseling (1991) |
| $h_4$ (m) | -80 | -80 | -80 | Wesseling (1991) |
| $\mathrm{K_{cb}}$ (-) | 1.15 | 1.15 | 1.15 | Allen et al. (1998) Table 17 Chapter 7 |
| $\mathrm{ET_o}$ ($\mathrm{md}^{-1}$) | 0.004 | 0.004 | 0.004 | Allen et al. (1998) Table 2 Chapter 1 |
| $\mathrm{f_c}$ (-) | 0.8 | 0.8 | 0.8 | Allen et al. (1998) Table 21 Chapter 7 |
| $A_{\mathcal{T}}$ ($\mathrm{m}^2$) | $\lvert\mathcal{A}_0\rvert = 0.088$ | $\mathrm{f_c}\lvert\mathcal{A}_{\mathrm{gbl}}\rvert = 1.35$ | $\mathrm{f_c}\lvert\mathcal{A}_{\mathrm{gbl}}\rvert = 1.35$ | N/A |
| $\mathrm{K_{cmax}}$ (-) | 1.30 | 1.30 | 1.30 | Allen et al. (1998) Chapter 7 |
| $\mathrm{f_w}$ (-) | 1 | 1 | 1 | Allen et al. (1998) Table 20 Chapter 7 |
| $\theta_{\mathrm{wp}}$ ($\mathrm{m}^3\mathrm{m}^{-3}$) | 0.15 | Multiple | 0.15 | Allen et al. (1998) Chapter 7 Table 19 |

| | | | | |
|---|---|---|---|---|
| $\theta_{\mathrm{fc}}$ ($\mathrm{m^3m^{-3}}$) | 0.29 | Multiple | 0.29 | Allen et al. (1998) Chapter 7 Table 19 |
| $\mathcal{T}_{3\mathrm{l}}$ ($\mathrm{md^{-1}}$) | $9.6 \times 10^{-4}$ | $9.6 \times 10^{-4}$ | $9.6 \times 10^{-4}$ | Cai et al. (2018) |
| $\mathcal{T}_{3\mathrm{h}}$ ($\mathrm{md^{-1}}$) | 0.0048 | 0.0048 | 0.0048 | Cai et al. (2018) |

**Table 2**: Model parameter values. The values with which the model in the main text was parametrised, for the various simulation scenarios.

| Soil type | Uniform constant initial pressure head profile $h_0$ in metres (m) |
|---|---|
| Clay | $h_0(x) = -5617.0$ for $x \in \Omega$ |
| Clay loam | $h_0(x) = -11.891$ for $x \in \Omega$ |
| Loam | $h_0(x) = -1.441$ for $x \in \Omega$ |
| Sandy clay loam | $h_0(x) = -1.132$ for $x \in \Omega$ |
| Silt | $h_0(x) = -6.185$ for $x \in \Omega$ |
| Silty clay loam | $h_0(x) = -77.245$ for $x \in \Omega$ |
| Silt loam | $h_0(x) = -5.0$ for $x \in \Omega$ |

**Table 3**: Initial pressure head profile used for each soil type in the investigation into the influence of preferential flow on water lifetime and uptake efficiency. These values ensure an initial uniform moisture content of $\theta(h_0(x)) = 0.214$ ($\mathrm{m^3m^{-3}}$), within there respective parametrisations of the soil water retention function $\theta$.
